# Impaired cholinergic modulation of striatal spiny projection neurons in repetitive behaviors associated with *TSHZ3*-linked autism spectrum disorder

**DOI:** 10.64898/2026.11.12.766099

**Authors:** Juliette Graniou, Xavier Caubit, Ahmed Fatmi, Abdel-Mouttalib Ouagazzal, Lydia Kerkerian-Le Goff, Laurent Fasano, Paolo Gubellini

## Abstract

Alterations in central cholinergic systems are implicated in autism spectrum disorder (ASD). We previously identified *TSHZ3* as an ASD-linked gene and provided evidence, from transgenic mouse models, supporting the view that striatal cholinergic interneurons (SCINs) are key players in the core ASD trait represented by repetitive behaviors (RBs). Indeed, conditional deletion of *Tshz3* in cholinergic neurons (*Chat-cKO* model) results in RBs associated with reduced SCIN firing activity and decreased striatal cholinergic tone. Here we tackled the resulting adaptations in the striatal circuitry underlying RBs in *Chat-cKO,* focusing on the two populations of striatal projection neurons (SPNs) that express dopamine D1 (D1R) or D2 (D2R) receptor. Using patch-clamp electrophysiological recordings, as well as behavioral, pharmacological and molecular approaches, we show reduced cholinergic transmission through the muscarinic M1 receptor onto SPNs, with no changes in the functionality or expression of this receptor. Notably, we demonstrate a loss of the differential excitability of D1R- *vs*. D2R-SPNs. Finally, acute *in vivo* treatment with the acetylcholinesterase inhibitor donepezil significantly alleviates RBs in mutant mice. These findings point to dysfunction of SCINs as a pathological and druggable mechanism underlying RBs in *Tshz3*-linked ASD, and to altered SPN excitability as a critical contributor in the resulting pathophysiological functioning of the striatum.

## INTRODUCTION

The striatum is the brain region containing the highest levels of acetylcholine, which is mainly released by striatal cholinergic interneurons (SCINs). Although SCINs represent only 1-2% of the whole striatal neuronal population, they have unique anatomofunctional features that confer them a pivotal modulatory role in the corticostriatal circuitry. Indeed, SCINs are characterized by extended dendritic and axonal arborization, allowing them to integrate afferent information from large striatal territories and to connect hundreds of neurons of the other subtypes (Lim *et al*., 2014). In addition, they have a spontaneous action potential firing activity both *in vivo* and *ex vivo* (Wilson *et al*., 1990; Bennett & Wilson, 1999; Apicella, 2017). This accounts for tonic striatal release of acetylcholine, which acts at both pre- and postsynaptic level via nicotinic and muscarinic receptors, regulating synaptic transmission/plasticity as well as neuronal excitability. Thereby, SCINs fine-tune information processing in the striatum and ultimately modulate its outflow via their action on spiny projection neurons (SPNs) (Calabresi *et al*., 2000; Pakhotin & Bracci, 2007; Bonsi *et al*., 2011; Abudukeyoumu *et al*., 2019; Assous, 2021). Both D1 receptor-expressing SPNs (D1R-SPNs) of the direct pathway and D2 receptor-expressing SPNs (D2R-SPNs) of the indirect pathway express M1 muscarinic receptors (M1Rs), while M4 receptors (M4Rs) are preferentially expressed by D1R-SPNs (Hersch *et al*., 1994; Ince *et al*., 1997; Santiago & Potter, 2001; Yan *et al*., 2001; Goldberg *et al*., 2012; Gonzales & Smith, 2015). Activation of M1Rs increases SPN excitability (Lv *et al*., 2017) by downregulating K_ir_2 channels (Shen *et al*., 2007), with a stronger effect in D2R-SPNs that are also more excitable than D1R-SPNs in basal conditions (Kreitzer & Malenka, 2007; Gertler *et al*., 2008). M4R activation decreases D1R-SPN excitability (Hernandez-Flores *et al*., 2015), although an opposite effect has been described (Figueroa *et al*., 2002). The cholinergic control is thus critical for the balanced activity between D1R- and D2R-SPNs required to enable action selection and maintain normal motor and cognitive functions, which accounts for the implication of SCIN dysfunction in a wide range of basal ganglia-related pathologies.

Clinical and animal studies point to molecular, neurodevelopmental and functional changes in cortical and striatal circuits among the possible neurobiological substrates of autism spectrum disorder (ASD) (Aliane *et al*., 2011; Delmonte *et al*., 2013; Shepherd, 2013; Fuccillo, 2016; Schuetze *et al*., 2016; Martos *et al*., 2017; Rapanelli *et al*., 2017a; Li & Pozzo-Miller, 2019; Thabault *et al*., 2022; Soghomonian, 2023; Evans *et al*., 2024) a neurodevelopmental disorder characterized by deficits in social interactions and repetitive behaviors (RBs) with restricted fields of interest (DSM-5, 2013). Abnormalities in the brain cholinergic system have been reported in patients with ASD (Rossignol & Frye, 2014; Eissa *et al*., 2018), and increasing evidence from animal models identify SCINs as possible players in this disorder (Karvat & Kimchi, 2014; Rapanelli *et al*., 2017b; Abudukeyoumu *et al*., 2019; Poppi *et al*., 2021). In this line, we previously identified *TSHZ3* (teashirt zinc-finger homeobox family member 3, also known as *ZFP537*) as a high confidence ASD-linked gene (Caubit *et al*., 2016; Feichtinger *et al*., 2022; Guin *et al*., 2025; Kesdiren *et al*., 2025) (see also https://omim.org/entry/617219 and https://gene.sfari.org/search/?search=tshz3). In the corticostriatal circuit, TSHZ3 is expressed in cortical projection neurons and, among the striatal neurons, in SCINs but not in SPNs. Using transgenic mouse models, we showed that conditional deletion (*cKO*) of *Tshz3* in cortical projection neurons results in social interaction deficits, while conditional deletion in neurons expressing the choline acetyltransferase gene (*Chat-cKO*) results in RBs (Caubit *et al*., 2022), revealing a dichotomy in the neural substrates of the two core ASD symptoms. Interestingly, *Chat-cKO* mice show an increased proportion of SCINs with low-frequency, irregular firing, resulting in an overall decrease in cholinergic activity, as assessed at presynaptic level by the reduced expression of the vesicular acetylcholine transporter (Molitor *et al*., 2026). Notably, RBs in these mice are likely associated to selective SCIN dysfunction, because *Tshz3* is expressed in all SCINs, while it is barely detectable or undetectable in the other cholinergic neurons of the brain and the brainstem, including those of the pedunculopontine and the laterodorsal tegmental nuclei that provide cholinergic inputs to the striatum (Dautan *et al*., 2014; Caubit *et al*., 2022). The *Chat-cKO* model is thus particularly useful and relevant to study SCIN-related dysfunctions in the striatal circuitry underlying RBs.

Here, we used this model to determine whether and how the decreased activity of SCINs impacts SPN excitability, and challenge the hypothesis that a treatment increasing cholinergic tone can alleviate *Tshz3* deletion-linked RBs. To do so, we studied M1R-mediated modulation of D1R- and D2R-SPN excitability in *Chat-cKO* mice *vs*. control, and tested whether an acute *in vivo* treatment with the acetylcholinesterase inhibitor donepezil can alleviate RBs in these mice.

## MATERIALS AND METHODS

### Ethics statement

All methods were performed in accordance with the relevant guidelines and regulations. Animal experimental procedures were in agreement with the recommendations of the European Communities Council Directive (2010/63/EU) and were approved by the French Ministry of Higher Education, Research and Space (ID number: APAFIS #45594-2023101113394773 v3).

### Mouse lines and genotyping

The *Tshz3*^flox/flox^, *Chat-Cre* and *Drd1a-tdTomato* (Jackson Laboratory #016204) mouse lines have been utilized and described previously (Mao *et al*., 1999; Madisen *et al*., 2010; Rossi *et al*., 2011; Chabbert *et al*., 2019; Caubit *et al*., 2022; Giua *et al*., 2024). Male heterozygous *Chat-Cre* were crossed with female *Tshz3*^flox/flox^ mice to generate *Chat-Cre*;*Tshz3*^flox/flox^ mice (“*Chat-cKO*”) with conditional KO (cKO) of *Tshz3* in cells expressing the enzyme choline acetyltransferase (*Chat*); *Cre*-negative littermates were used as control. Triple transgenic *Chat-Cre*;*Tshz3*^flox/flox^;*Drd1a-tdTomato* mice (“*tdTom-Chat-cKO*”) and their relative *Cre*-negative littermate controls were used to identify D1R-SPNs via the red fluorescence of tdTomato, a modified version of the DsRed fluorescent protein. Animals were genotyped as described previously (Chabbert *et al*., 2019; Caubit *et al*., 2022). Only male mice were used, and no randomization was performed. Animals were excluded from the analyses if their score in the marble burying test (MBT), or in the D1/D2 ratio (see Results), was identified as an outlier (see Statistical analysis).

### Electrophysiology

Electrophysiological data were obtained from 18 *Chat-cKO* mice and 20 *Cre*-negative littermate controls, and from 29 *tdTom-Chat-cKO* mice and 32 *Cre*-negative littermate controls. All mice were male, aged between 21 and 28 days. Procedures were similar to those described previously (Caubit *et al*., 2016; Chabbert *et al*., 2019; Caubit *et al*., 2022). Briefly, acute coronal brain slices (250 µm-thick) were cut using a S1000 Vibratome (Leica Biosystems, Nussloch, Germany) in ice-cold solution containing (in mM): 110 choline, 2.5 KCl, 1.25 NaH_2_PO_4_, 7 MgCl_2_, 0.5 CaCl_2_, 25 NaHCO_3_ and 7 glucose, pH 7.4. Slices were kept at room temperature in oxygenated artificial cerebrospinal fluid (ACSF), whose composition was (in mM): 126 NaCl, 2.5 KCl, 1.2 MgCl_2_, 1.2 NaH_2_PO_4_, 2.4 CaCl_2_, 11 glucose and 25 NaHCO_3_, pH 7.4. Electrophysiological recordings were performed in ACSF at 34-35 °C and flowing at ∼2 ml/min. Striatal SPNs were identified by infrared video microscopy and by their electrophysiological properties, namely hyperpolarized resting membrane potential (RMP) (around −75 mV), low input resistance (60-100 MΩ) and slowly-developing depolarization preceding the emission of action potentials (Jiang & North, 1991; Cepeda *et al*., 1994; Kawaguchi, 1997). In experiments using *tdTom-Chat-cKO* mice, D1R-SPNs and putative D2R-SPNs were identified due to the presence or the absence of tdTomato fluorescence, respectively. SPNs were recorded (sampling rate 10 kHz) in whole-cell configuration using patch-clamp borosilicate micropipettes (4-5 MΩ) filled with an internal solution containing (in mM): 125 K-gluconate, 10 NaCl, 1 CaCl_2_, 2 MgCl_2_, 0.5 BAPTA, 19 HEPES, 0.3 Na-GTP and 1 Mg-ATP, pH 7.3. The recording pipette was moved by a MP-225 electric micromanipulator (Sutter Instrument, Novato, CA, USA). Electrophysiological data were recorded and digitized by an AxoPatch 200B amplifier, a Digidata 1550B4 signal converter and pClamp 10.7 software (Molecular Devices, Wokingham, UK). Analysis of electrophysiological parameters was performed blinded to the genotype. Current-voltage (I-V) relationship was obtained by injecting hyperpolarizing and depolarizing current steps (ΔI = 50 pA, 500 ms), and input resistance was calculated for each recorded neuron by linear regression analysis, i.e., as the slope of the linear best fit of its I-V relationship. SPN excitability was studied by injecting 500 ms current steps with 100 pA increments; in particular, a +300 pA step was used as reference because it was the minimal current intensity capable of systematically triggering action potentials. The rheobase was measured as the minimal current capable of triggering at least one action potential, using 500 ms current steps from 0 to +400 pA with 10 pA increments. Drugs were applied in the ACSF at the concentration and duration indicated in the Figures and/or the Results. VU 0357017 hydrochloride and McN-A 343 were purchased from Tocris Cookson Ltd. (Bristol, UK), tropicamide and donepezil from Merck KGaA (Darmstadt, Germany).

### Marble burying test

24 *Chat-cKO* mice and 32 littermate controls underwent the marble burying test (MBT). The protocol used here has been previously detailed and used to quantify stereotyped behavior (Caubit *et al*., 2016; Caubit *et al*., 2022), and has been validated to model ASD-related RBs in mouse (Njung’e & Handley, 1991; Thomas *et al*., 2009; Roubertoux *et al*., 2020). It consists of placing a mouse in a 40 x 18 cm cage with 5 cm-thick litter and containing 20 marbles (9 mm diameter) on the surface of the litter. At the end of a 15-minute session, the degree at which each marble is buried by the mouse is scored 2, 1 and 0 when completely buried, buried but still partially visible and not buried, respectively, and values for the 20 marbles are summed (MBT score thus ranges from 0 to 40). Before performing the MBT, each mouse was individually placed into the cage containing only the litter and no marble for a 10-minute habituation period. Then, the mouse was removed, the 20 marbles were placed on the litter in 4 lines of 5 marbles, and it was placed back in the cage to perform the MBT.

### Donepezil treatment

Donepezil (Dpz) was diluted in vehicle (0.9% NaCl) at a concentration of 0.2 mg/mL (528 µM) in order to obtain a dose of 1 mg/kg by injecting (i.p.) 5 µL of solution per gram of mouse. Acute Dpz treatment *vs*. vehicle (NaCl) was tested on 16 *Chat-cKO* mice and 21 littermate controls, which were subdivided in 4 age-matched cohorts of 4-5 mice each. They were aged 20 weeks at the time of the 1^st^ MBT session. Each cohort was split in two sub-cohorts, each containing a similar number of control and *Chat-cKO* mice: one sub-cohort received Dpz and the other NaCl, and after 30 min each mouse underwent the MBT. Two weeks later, mice previously injected with Dpz received NaCl and *vice versa*, and 30 min later the MBT was performed. The MBT scores were then pooled for each mouse in NaCl and Dpz condition, thus allowing NaCl *vs*. Dpz paired comparisons for both genotypes.

A supplementary cohort of 8 *Chat-cKO* mice and 11 littermate controls underwent two times the MBT at 2-week interval without receiving any treatment (naïve condition), in order to validate the stability of the score between the two consecutive sessions and the repeatability/reproducibility of this behavioral assessment used to test the effect of NaCl *vs*. Dpz (see Fig. 6A, B).

### RT-qPCR

Experiments were performed by an investigator blinded to genotype on samples of acutely dissected striata from 10 *Chat-cKO* mice and 5 littermate controls, aged 25-30 days. Total RNA was extracted using Direct-zol RNA Microprep Kit (Zymo Research, CA, USA) and first-strand cDNA was synthesized with SuperScript™ IV VILO™ Master Mix (Thermo Fisher Scientific Inc., Waltham, MA, USA). Real-time quantitative polymerase chain reaction (RT-qPCR) was carried out on a CFX96 RT PCR Detection System (Bio-Rad Laboratories Inc., Hercules, CA, USA) using PowerTrack™ SYBR Green Master Mix (Thermo Fisher Scientific Inc.). Thermal cycling conditions consisted of 40 cycles at 95 °C for 15 s and at 60 °C for 60 s. All reactions were performed in triplicate. Relative expression levels of mouse *Chrm1* (type 1 muscarinic receptor, M1R) and *Chrm4* (type 4 muscarinic receptor, M4R) mRNAs were calculated using the 2^−ΔΔCt^ method, with *Hprt* (hypoxanthine guanine phosphoribosyl transferase) as the reference housekeeping gene for normalization and the mean value of the control group as the calibrator. The stability of *Hprt* expression between control and *Chat-cKO* mice was assessed prior to its use as a reference gene for normalization. No significant genotype-dependent variation was observed. The specificity of the RT-qPCR amplification products was assessed by melting curve analysis, which revealed a single distinct melting peak for each primer pair, with no evidence of non-specific amplification or primer-dimer formation. Amplification efficiencies were experimentally determined using five-point standard dilution series of cDNA (10⁰ to 10⁻⁴), with the initial 1:3 cDNA dilution defined as 10⁰. Only dilution points within the linear amplification range were included in the standard curve regression analysis. All primer pairs (*Chrm1*, *Chrm4* and *Hprt*) exhibited amplification efficiencies close to 100%, with comparable slopes across the standard curves, supporting the use of the 2⁻ΔΔCt method for relative gene expression quantification.

The primer sequences were:

Fw-*Chrm1:* 5’-GGTTGGTTTCCTTCGTTCTCTGG-3’ (23b);

Rev-*Chrm1:* 5’-GGAGAGGAACTGGATGTAGCAC-3’ (22b);

Fw-*Chrm4:* 5’-CCTTCTACCTGCCTGTGGTCAT-3’ (22b);

Rev-*Chrm4:* 5’-TGAGGAAAGCCAGAGTCTTGGC-3’ (22b);

Fw-*Hprt:* 5’-CTGGTGAAAAGGACCTCTCGAAG-3’ (23b);

Rev-*Hprt:* 5’-CCAGTTTCACTAATGACACAAACG-3’ (24b).

### Statistical analysis

The choice of sample sizes was based on previous experience and by the results of the on-line tool BiostaTGV (https://biostatgv.sentiweb.fr/?module=etudes/sujets#, developed by the Pierre Louis Institute, the INSERM and the Sorbonne University), which calculates the sample size based on the expected standard deviation and the required alpha risk and power. Statistical analysis was performed using Prism 8.4.3 (GraphPad Software, Boston, MA, USA). Data sets were tested for normality by the D’Agostino & Pearson’s and the Shapiro-Wilk tests. Outliers were identified by the ROUT method (Motulsky & Brown, 2006) and excluded. Data sets were compared using parametric tests if they all showed a normal distribution; in this case, they are expressed as mean ± standard error of the mean (SEM). Otherwise, if at least one data set did not show a normal distribution, they were analyzed by non-parametric tests and expressed as median ± interquartile range (IQR) or median followed by 1^st^ and 3^rd^ quartiles. In some cases, single values are shown individually as dots. One- and two-sample Student’s *t*-test (unpaired and paired), Mann-Whitney test and Wilcoxon test were used for comparing two data sets. ANOVA, eventually followed by uncorrected Fisher’s least significant difference (LSD) post-test, and Kruskal-Wallis test, were used to compare 3 or more data sets. The significance threshold was set at *P* < 0.05.

Bimodal *vs*. unimodal distribution and bimodality coefficient (BC) of SPN firing frequency were calculated using RStudio 4.2.1 free software (Posit Software, Boston, MA, USA; https://www.r-project.org/) and the “mclust” package (Scrucca *et al*., 2016) for model-based clustering, classification, and density estimation (https://cran.r-project.org/web/packages/mclust/vignettes/mclust.html). The BC ranges from 0 to 1, where BC > 0.555 and BC < 0.555 correspond, respectively, to a bimodal and unimodal distribution (Freeman & Dale, 2013; Pfister *et al*., 2013).

The statistical tests used, *P* values, sample sizes [as number (n) of SPNs or mice] and the way data are represented are indicated in the Methods, Results and/or Figures.

## RESULTS

### Cholinergic transmission onto SPNs through M1R

We previously showed that *Chat-cKO* mice have reduced SCINs firing frequency resulting in decreased cholinergic activity (Molitor *et al*., 2026), but the functional consequences of these changes at post-synaptic level still need to be characterized to understand the substrates of RBs. For instance, acetylcholine-mediated modulation of SPN excitability could be impaired, and/or the functionality and expression of the muscarinic receptors expressed by SPNs could be modified.

To challenge these hypotheses, during slice patch-clamp recordings we applied the M1R-selective positive allosteric modulator (PAM) VU 0357017, which should provide an excitatory effect proportional to the basal activation level of M1Rs (Lv *et al*., 2017). We therefore expected this PAM to increase SPN firing frequency in control mice, and to have a lower or no effect in *Chat-cKO* due to their decreased cholinergic tone in the striatum. Accordingly, the application of VU 0357017 (10 µM, 15 min) on slices of control mice significantly increased SPN firing frequency, but had no effect in *Chat-cKO* (Fig. 1A and B). This result provides functional evidence for decreased striatal cholinergic transmission through M1 receptors onto SPNs in *Chat-cKO* mice, which could impact SPNs excitability and thereby striatal circuitry functioning and output.

**Figure 1.**
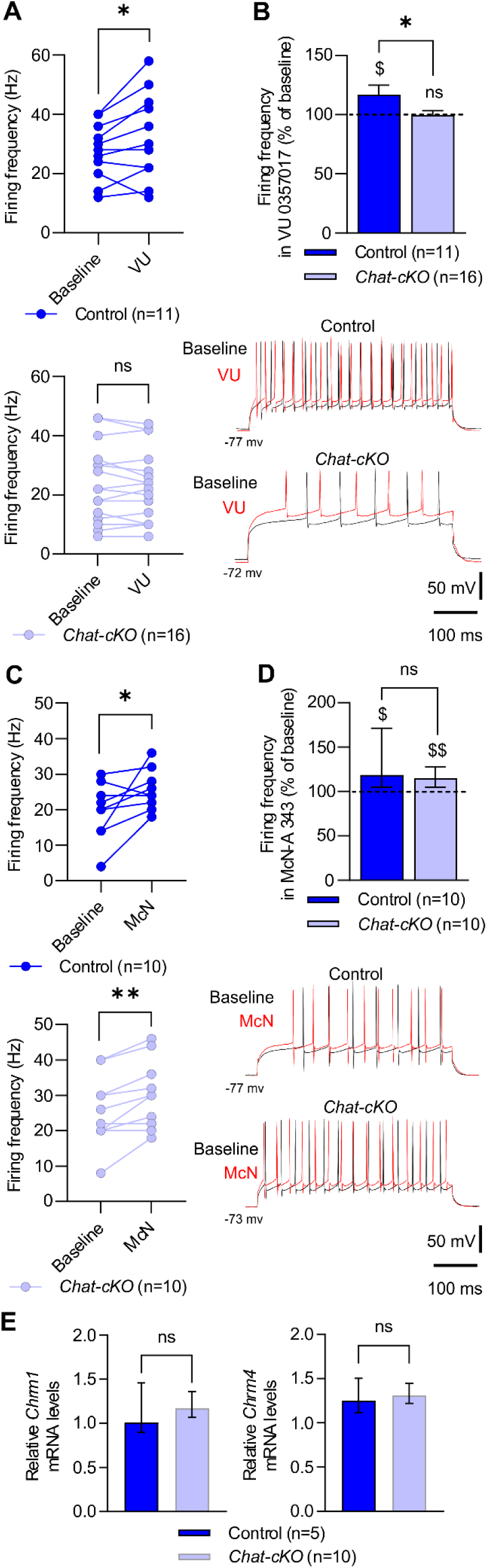
M1R-mediated SPN excitability, and expression of M1R and M4R mRNAs. (**A**) The M1R-selective PAM VU 0357017 (VU; 10 µM, 15 min) significantly increased SPN excitability in control (upper graph) but not in *Chat-cKO* (lower graph) mice (paired Student’s *t*-test: \**P* < 0.05, ns = non-significant). (**B**) The excitatory effect of VU 0357017 was significant only in control mice, and this effect was significantly different between the two genotypes (Student’s *t*-test: \**P* < 0.05; paired Student’s *t*-test: ^$^*P* < 0.05, ns = non-significant *vs*. baseline). (**C**) The M1R-selective agonist McN-A 343 (McN; 10 µM, 5 min) provided a significant increase in SPN excitability in both control (upper graph) and *Chat-cKO* (lower graph) mice (paired Student’s *t*-test: \**P* < 0.05, \*\**P* < 0.01). (**D**) The excitatory effect of McN-A 343 was significant and similar in control and *Chat-cKO* mice (Student’s *t*-test: ns = non-significant; Wilcoxon signed rank test: ^$^*P* < 0.05, ^$$^*P* < 0.01 *vs*. baseline). Traces show SPN firing from control and *Chat-cKO* mice before (black) and after (red) the application of the drugs; note the variability of firing frequency among different SPNs, also visible on the graphs in A and C. (**E**) The relative striatal expression of M1R (*Chrm1*) and M4R (*Chrm4*) mRNAs, measured by RT-qPCR, is similar in control and *Chat-cKO* mice (Mann-Whitney test: ns = non-significant). Data in A and C are expressed as individual values; in B as mean + SEM; in D and E as median ± IQR; “n” in A-D refer to the number of recorded SPNs, in E to the number of animals.

We then studied whether and how M1R function is altered in *Chat-cKO* compared to control mice, knowing that the pharmacological activation of this receptor increases SPN firing frequency (Lv *et al*., 2017). For this, we tested the effect of the selective M1R agonist McN-A 343 (Calabresi *et al*., 1998) by measuring the frequency of SPN action potential firing triggered by the injection of depolarizing current steps (300 pA, 500 ms), before and after drug application. McN-A 343 (10 µM, 5 min) significantly increased SPN firing frequency in both control and *Chat-cKO* mice (Fig. 1C), with a similar effect in both genotypes (Fig. 1D). Finally, we used RT-qPCR to measure the levels of *Chrm1* and *Chrm4* mRNAs, which encode the M1R and M4R, respectively, in the striatum of control and *Chat-cKO* mice. This approach did not reveal statistically significant differences in striatal *Chrm1* and *Chrm4* mRNA levels between control and *Chat-cKO* mice under the experimental conditions tested (Fig. 1E).

These results suggest that, despite a reduced basal activity of M1R in *Chat-cKO* mice likely due to the lower striatal cholinergic tone, both the function and the expression of this receptor are preserved.

### Electrophysiological properties of SPNs

These changes in acetylcholine receptor activation could affect the membrane properties of SPNs and, ultimately, modify their excitability (Oldenburg & Ding, 2011). We thus studied SPN current-voltage (I-V) relationship, which was unchanged between control and *Chat-cKO* mice (Fig. 2A), and provided similar input resistance values (90.08 ± 3.54 and 90.12 ± 3.83 MΩ, respectively; Mann-Whitney test: *P* = 0.596; values are expressed as mean ± SEM). The resting membrane potential (RMP) was also similar in SPNs from control and *Chat-cKO* mice (Fig. 2B), while the rheobase was slightly higher in *Tshz3*-mutant mice, but this difference was not significant (Fig. 2C). These results demonstrate that these membrane properties are not significantly impacted in *Chat-cKO* mice.

**Figure 2.**
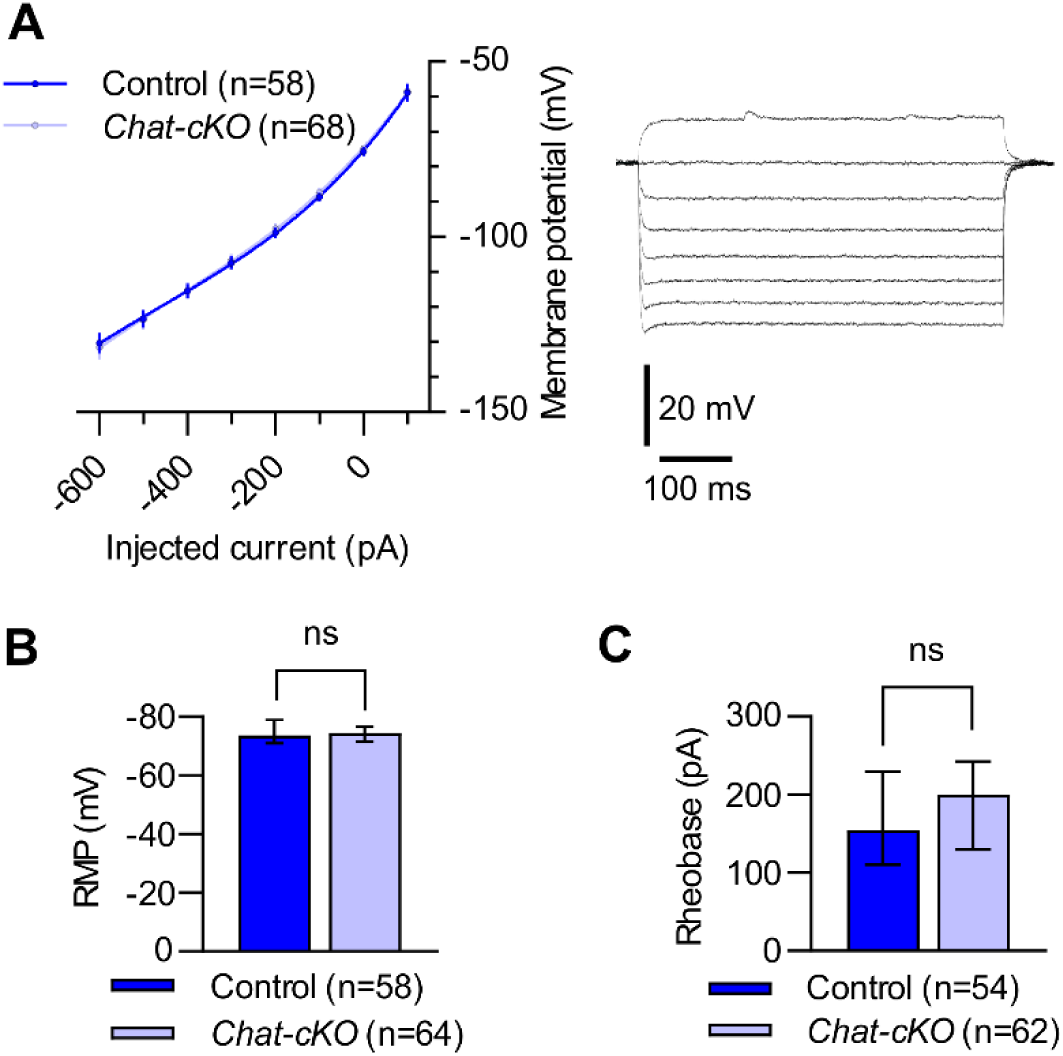
SPN membrane properties. (**A**) I-V relationships (left graph) of control and *Chat-cKO* SPNs largely overlap. Note the inward rectification typical of these neurons (curves are 3^rd^ order polynomial fits). Traces (right) show the membrane responses of a control SPN to hyperpolarizing and depolarizing current steps from −600 to +100 pA, with 100 pA increments. (**B**) Resting membrane potential (RMP) and (**C**) rheobase are similar in SPNs from control and *Chat-cKO* mice (Mann-Whitney test: ns = non-significant). Data in A are expressed as mean ± SEM; some error bars are smaller than the dots representing the average values, which overlap. Data in B and C are expressed as median ± IQR; “n” refer to the number of recorded SPNs.

SPN excitability, again measured as the firing frequency in response to a depolarizing current step, was also similar between control and *Chat-cKO* mice (Fig. 3A). Intriguingly, however, the distribution of the frequency values was different between the SPNs of the two genotypes (Fig. 3B and C). Indeed, clustering analysis (see Methods) identified 2 distinct clusters in control SPNs while only 1 in *Chat-cKO,* and provided, respectively, bimodality coefficients of 0.603 and 0.489 that clearly correspond to bimodal and unimodal distribution. The presence of two clusters in control mice is consistent with the literature data showing that the two populations of SPNs have differential excitability, with D2R-SPNs being more excitable than D1R-SPNs (Kreitzer & Malenka, 2007; Gertler *et al*., 2008). The unimodal distribution in *Chat-cKO* mice may thus reflect a loss of this asymmetry.

**Figure 3.**
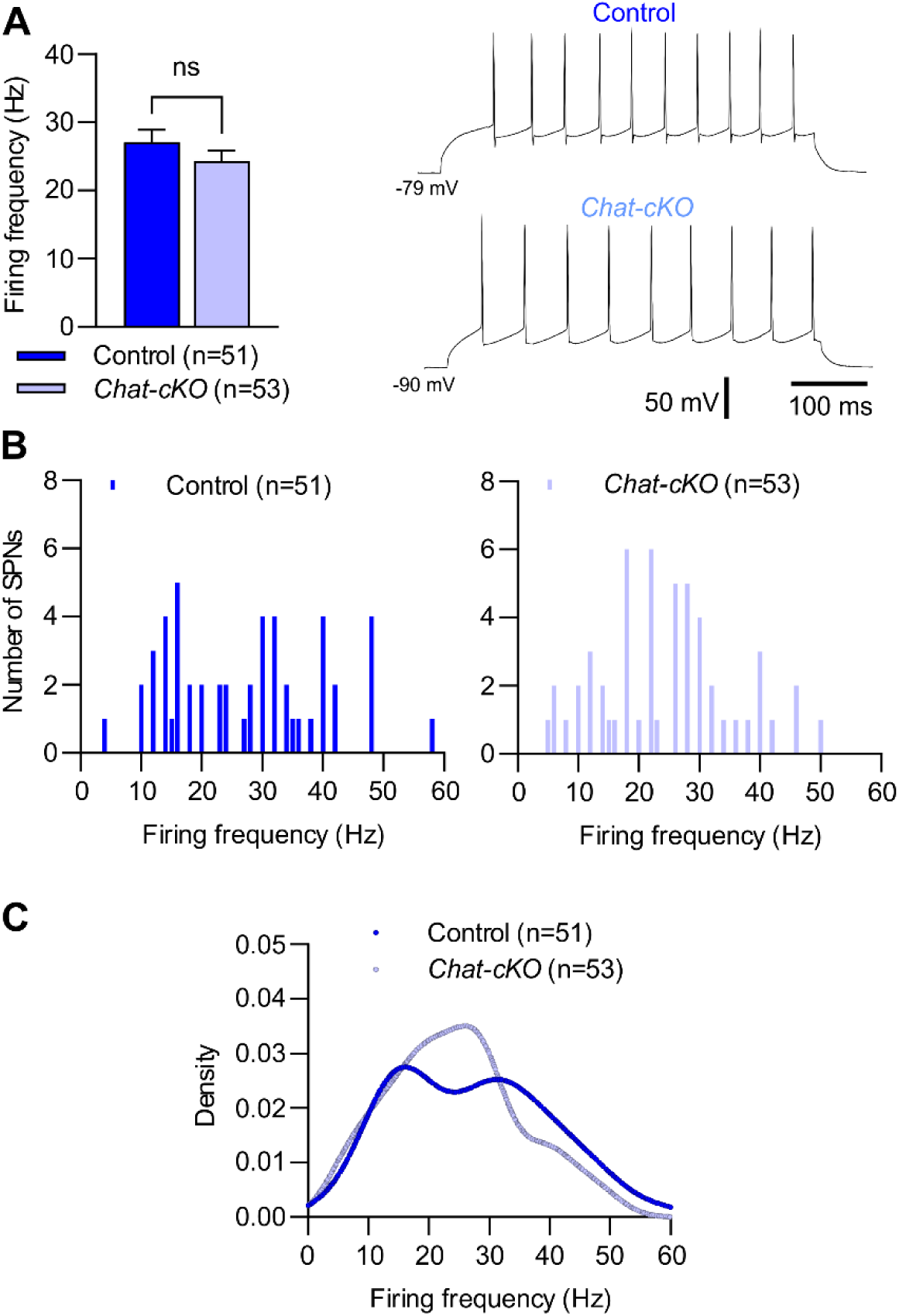
SPN excitability. (**A**) The average SPN firing frequency in response to a +300 pA, 500 ms current step (left graph) is similar in control *vs*. *Chat-cKO* mice (Student’s *t*-test test: ns = non-significant). Traces (right) show the responses of a control and a *Chat-cKO* SPN to a +300 pA depolarizing current step. (**B**) Distribution (bin width = 1) of SPN firing frequency values in control (left) and *Chat-cKO* (right) mice. (**C**) The same data as in B expressed as superposed kernel density plots, clearly showing the bimodal *vs*. unimodal distribution of control and *Chat-cKO* SPNs, respectively. Data in A are expressed as mean + SEM; “n” refer to the number of recorded SPNs.

Overall, these findings indicate that the reduced striatal cholinergic tone resulting from *Tshz3* deletion in SCINs leads to a homogenization of SPN excitability, which might profoundly affect striatal function.

### Electrophysiological properties of D1R-SPNs and D2R-SPNs

The unimodal/homogeneous distribution of SPN excitability found in *Chat-cKO* mice is intriguing and calls for a thorough study allowing the differentiation between D1R- and D2R-SPNs. To achieve this, we generated triple transgenic *tdTom-Chat-cKO* mice, and their littermate controls, in which D1R-SPNs were identified by their expression of the tdTomato red fluorescent protein, while SPNs not exhibiting red fluorescence were categorised as D2R-SPNs (see Methods).

First, we characterized the membrane properties of D1R- and D2R-SPNs in both control and *tdTom-Chat-cKO* mice. Overall, no significant differences were observed between genotypes or between the two SPN populations concerning RMP (Fig. 4A) and input resistance [control: D1R-SPNs (n=30) 64.9 MΩ (49.8;78.9), D2R-SPNs (n=19) 72.0 MΩ (56.8;91.8); *tdTom-Chat-cKO*: D1R-SPNs (n=21) 60.4 MΩ (55.5;81.9), D2R-SPNs (n=25) 67.6 MΩ (55.5;90.1); Kruskal-Wallis test: *P* = 0.479; values are expressed as medians followed by the 1^st^ and 3^rd^ quartiles]. In contrast, in control mice, the rheobase of D2R-SPNs was significantly lower than that of D1R-SPNs, which is consistent with their higher excitability (Kreitzer & Malenka, 2007; Gertler *et al*., 2008). Interestingly, this difference was absent in *tdTom-Chat-cKO* mice, where D1R- and D2R-SPNs showed similar rheobase values (Fig. 4B).

**Figure 4.**
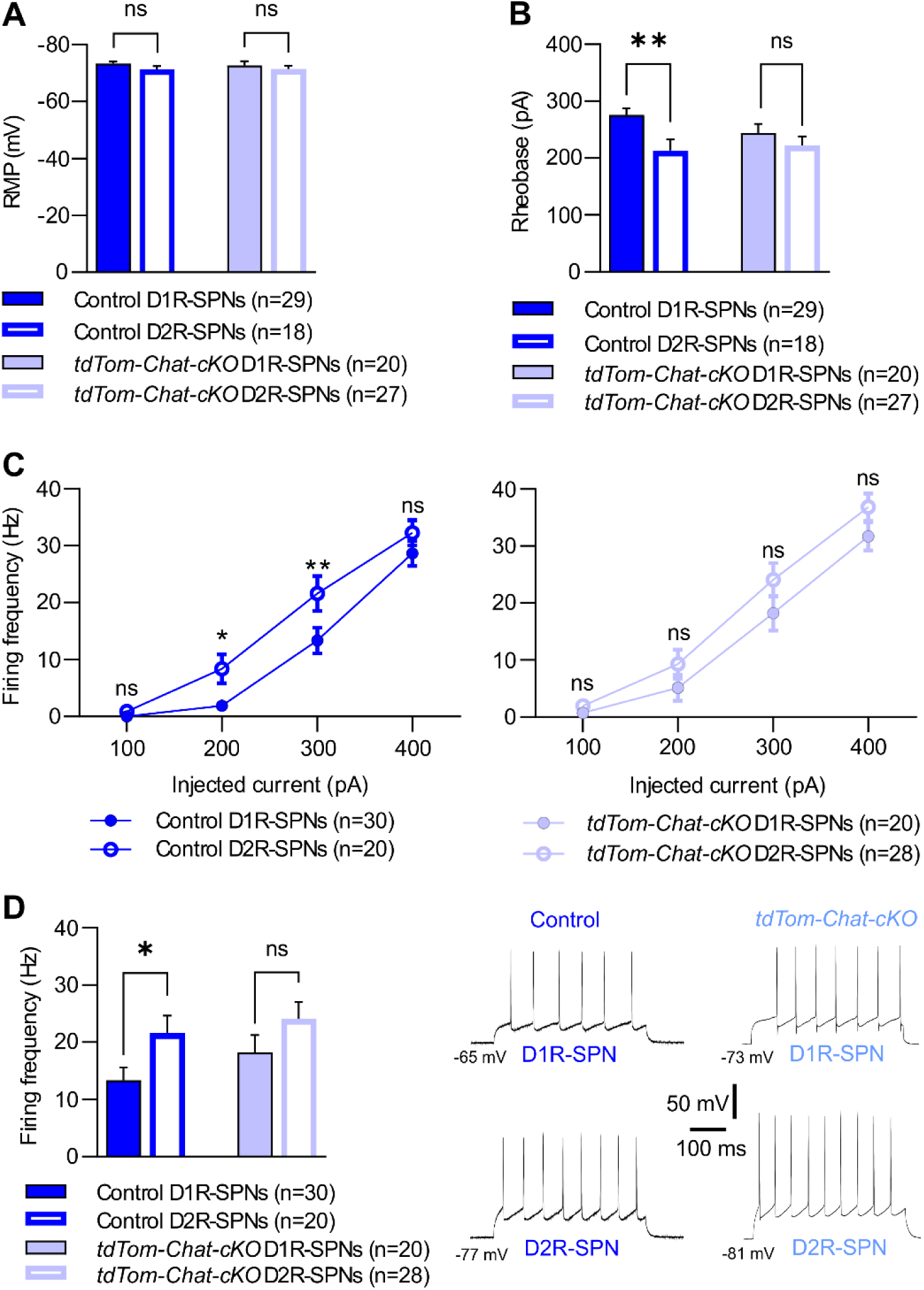
D1R-SPN *vs*. D2R-SPN excitability. (**A**) The resting membrane potential (RMP) is similar between D1R-SPNs and D2R-SPNs in both control and *tdTom-Chat-cKO* mice, as well as between control and *tdTom-Chat-cKO* SPNs of the same population (ANOVA: *F*(3,90) = 0.8145, *P* = 0.489). (**B**) The rheobase of control D2R-SPNs is significantly lower than D1R-SPNs, while in *tdTom-Chat-cKO* mice the two SPN populations have similar rheobase values (ANOVA: *F*(3,90) = 3.532, *P* = 0.018; Fisher’s LSD post-test: \*\**P* < 0.01, ns = non-significant; for each SPN subtype, comparison of control *vs*. *tdTom-Chat-cKO* was non-significant). (**C**) Responses of D1R-SPNs and D2R-SPNs from control (left) and *tdTom-Chat-cKO* (right) mice to progressively increasing current steps clearly show that D2R-SPNs are more excitable than D1R-SPNs in control mice, while their excitability is similar in *tdTom-Chat-cKO* (Student’s *t*-test: \**P* < 0.05, \*\**P* < 0.01, ns = non-significant). (**D**) The average D2R-SPN firing frequency in response to +300 pA current steps (same as in C) is significantly higher than D1R-SPNs in control mice, while in *Chat-cKO* the difference is not significant (ANOVA: *F*(3,94) = 3.093, *P* = 0.0307; Fisher’s LSD post-test: \**P* < 0.05, ns = non-significant; for each SPN subtype, comparison of control *vs*. *tdTom-Chat-cKO* was non-significant). Traces show the responses of D1R-SPNs and D2R-SPNs from control and *tdTom-Chat-cKO* mice. Data are expressed as mean ± SEM; “n” refer to the number of recorded SPNs.

We then studied SPN firing frequency in response to current steps of increasing intensity and, in agreement with the literature, control D2R-SPNs showed significantly higher excitability than D1R-SPNs (Fig. 4C). Conversely, in *tdTom-Chat-cKO* mice, no significant difference was found between the two SPN populations (Fig. 4C). The average responses to +300 pA current steps are graphed in Fig. 4D, illustrating the loss of the differential D1R- *vs*. D2R-SPN excitability in *Tshz3*-mutant mice, with a slightly increased excitability of D1R-SPNs.

We also studied these selective SPN changes by calculating the ratio between the average firing frequency of D1R-over D2R-SPNs (D1R/D2R) for each mouse, in order to minimize the variability of this parameter between different animals, cohorts and experimental sessions. Analysis by one-sample *t*-test *vs*. a value of 1 clearly shows that the D1R/D2R ratio is significantly smaller than 1 in control mice, confirming that the firing frequency of D2R-SPN is higher than that of D1R-SPNs, while *Tshz3*-mutant mice show a ratio close to 1, indicating similar excitability of the two SPN populations (Fig. 5). Accordingly, the D1R/D2R ratio is also significantly lower in control *vs*. *tdTom-Chat-cKO* mice (Fig. 5). These data comply with the change in the firing frequency distribution of the global population of SPNs from bimodal to unimodal in control *vs*. *Chat-cKO* mice shown in Fig. 3C, confirming the loss of the differential excitability of the two populations of SPNs.

**Figure 5.**
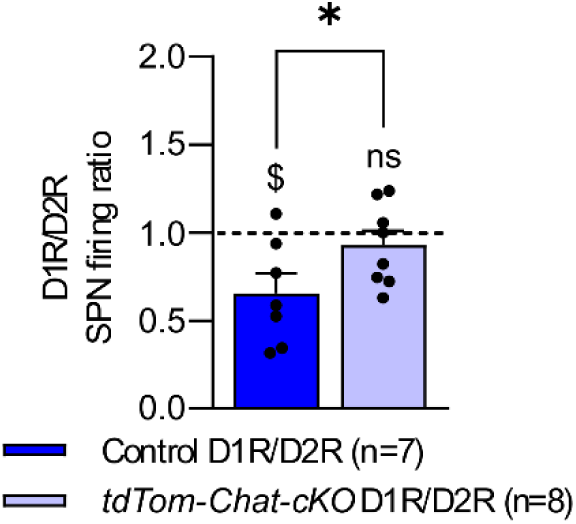
Ratio between the average firing frequency of D1R- and D2R-SPNs (D1R/D2R) per mouse. The D1R/D2R ratio is significantly lower than 1 in control mice, consistent with the higher excitability of D2R-SPNs, while it is close to 1 in *tdTom-Chat-cKO* (one-sample *t*-test *vs*. D1R/D2R = 1: ^$^*P* < 0.05, ns = non-significant), with a significant difference between the two genotypes (Student’s *t*-test with Welch’s correction: \**P* < 0.05). Data are expressed as mean + SEM; “n” refer to the number of mice in which SPNs were recorded; for each mouse, the D1R/D2R ratio is obtained from 1 to 4 SPNs of each subtype.

Overall, these electrophysiological findings indicate that the reduced striatal cholinergic tone in *Tshz3*-mutant mice does not drastically affect SPN membrane properties, but rather homogenizes D1R- and D2R-SPN excitability, possibly due to increased D1R-SPN excitability. This could affect the way these neurons differentially respond to and integrate their synaptic inputs, and consequently the balance between the direct and indirect output pathways of the striatum.

### Repetitive behaviors and cholinergic treatment with donepezil

The data presented above and in our recent paper (Molitor *et al*., 2026) strongly support the idea that reduced striatal cholinergic tone in *Chat-cKO* mice plays a crucial role in RBs by untuning SPN excitability. Therefore, we tested the hypothesis that enhancing cholinergic transmission could improve this pathological behavior. To this end, we assessed the effects of an acute treatment of *Chat-cKO* and control mice with the acetylcholinesterase inhibitor donepezil (Dpz) using the marble burying test (MBT). First, we tested whether mice repeating the MBT twice would provide stable and reproducible scores, and whether the scores differ between animals of the two genotypes. As shown in Fig. 6A, there was no significant difference in the scores between the 1^st^ and the 2^nd^ MBT session at 2-week interval, in both control and *Chat-cKO* mice. Averaging per mouse the two scores revealed a significantly higher score in *Chat-cKO* mice compared to control (Fig. 6B). These results confirm our previous findings obtained with that Chat-cKO mice exhibit RBs as evidenced with the MBT (Caubit *et al*., 2022) and show that this test provides stable results when repeated twice, thus validating it for comparing the effect of repeated Dpz *vs*. saline treatment in the same animal as for the next experiment.

**Figure 6.**
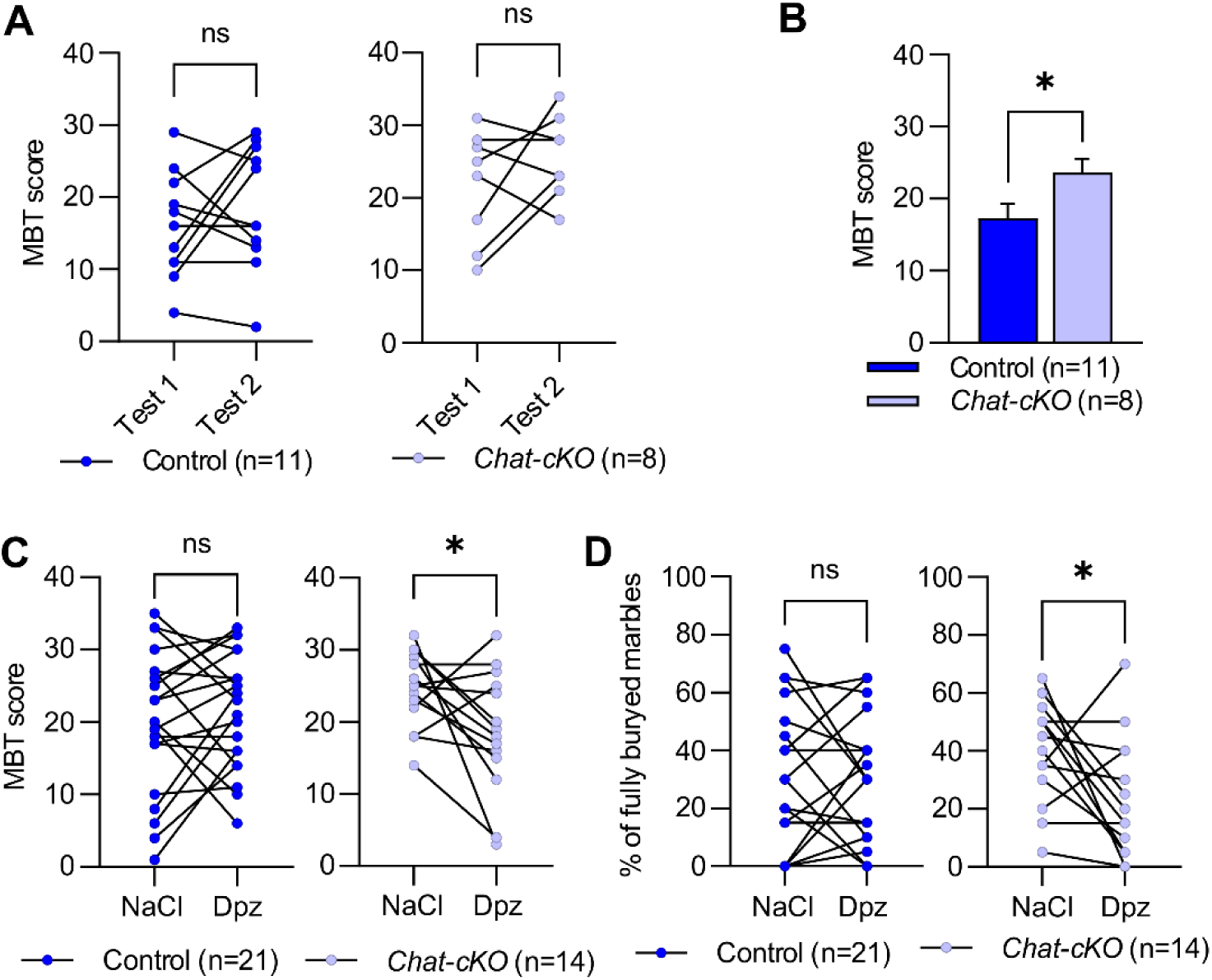
RBs and cholinergic treatment. (**A**) The MBT provided stable scores between 2 sessions (Test 1 and 2) at 2-week interval, in both control and *Chat-cKO* naïve mice (paired Student’s *t*-test: ns = non-significant). (**B**) Averaging the data obtained per mouse in Test 1 and 2 (as in A) shows a significantly higher MBT score in mutant mice (Student’s *t*-test: \**P* < 0.05). (**C**, **D**) Treatment with the acetylcholinesterase inhibitor donepezil (Dpz, 1 mg/kg i.p.) significantly reduced RBs in *Chat-cKO* mice *vs*. vehicle (NaCl 0.9%) either in terms of MBT score (**B**) or percent of fully buried marbles (**C**), while it had no significant effect in control mice (paired Student’s *t*-test: \**P* < 0.05, ns = non-significant). Note that, in D, several points are superposed at values close or equal to zero. Data in A, C and D are expressed as individual mouse values; data in B are expressed as mean + SEM; “n” refer to the number of tested animals.

Using different mouse cohorts (see Methods), we then assessed the outcome of Dpz treatment (1 mg/kg, i.p.) *vs*. vehicle (0.09% NaCl). Interestingly, Dpz significantly reduced RBs in *Chat-cKO* mice measured as MBT score (Fig. 6C) as well as percentage of fully-buried marbles (Fig. 6D), while it had no significant effect in controls.

Overall, these data clearly demonstrate that enhancing the cholinergic tone with acute donepezil treatment significantly alleviates the RBs that characterize *Chat-cKO* mice. This supports the critical role of SCINs and striatal acetylcholine in this ASD-related pathological behavior caused by conditional *Tshz3* deletion.

## DISCUSSION

Although a decreased cholinergic activity in the striatum seems clearly involved in RBs and ASD, the implicated striatal circuitry components remain somehow elusive. The striatum is a key regulatory center integrating, filtering and transforming the information arising from virtually all the cortical areas, the intralaminar and ventrolateral thalamic nuclei, and from other basal ganglia structures such as the substantia nigra pars compacta and the external globus pallidus (DeLong & Wichmann, 2007). The information processed in the striatum is then transmitted to the output structures of the basal ganglia – the internal globus pallidus (GPi) and the substantia nigra pars reticulata (SNr) – and then back to the cerebral cortex via the thalamus, forming the three (motor, associative and limbic) cortico-basal ganglia-thalamo-cortical “loops” (Macpherson & Hikida, 2019). The striatal output to the GPi/SNr is constituted of the “direct” pathway originating from D1R-SPNs, and the “indirect” pathway originating from D2R-SPNs and involving the external globus pallidus and the subthalamic nucleus. These two pathways exert an opposite control on the selection of motor programs: namely, the activation of the direct pathway is considered to initiate a given motor program, while the concomitant activation of the indirect pathway inhibits concurrent or inappropriate motor programs. A subtle balance between the activity of D1R- and D2R-SPNs is thus crucial for the selection of relevant actions and the suppression of unwanted movements (Calabresi *et al*., 2014; Hashemiyoon *et al*., 2017; Obeso *et al*., 2017; Gerfen, 2022). It is long known that acetylcholine, released in the striatum mainly by SCINs, has a powerful modulatory effect on both synaptic transmission/plasticity and SPN excitability and, ultimately, in the fine balance of striatal output (Calabresi *et al*., 2000; Figueroa *et al*., 2002; Pakhotin & Bracci, 2007; Bonsi *et al*., 2011; Gonzales & Smith, 2015; Lv *et al*., 2017; Abudukeyoumu *et al*., 2019; Assous, 2021). Indeed, SCINs play a crucial role in the behavioral flexibility required for switching and selecting the appropriate behaviors in response to changing environmental conditions (Apicella, 2017). In this context, we focused here on the possible consequences of the decreased striatal cholinergic tone in *Chat-cKO* mice on the electrophysiological properties of SPNs. First, we assessed its functional outcome showing, as expected, that the effect of the M1R PAM VU 0357017 is significantly reduced, suggesting a lower basal activation of this receptor. Secondly, we studied the function of M1Rs by testing the effect of the M1R agonist McN-A 343 on SPN excitability, and the expression of M1Rs and M4Rs by measuring their mRNA levels, and our results clearly show that they are not affected by *Tshz3* mutation. Therefore, the lower cholinergic tone in the striatum of *Chat-cKO* mice results in a lower basal activation of M1Rs, but does not affect their sensitivity and expression. Although these data are conclusive, it cannot be excluded that these electrophysiological and neuropharmacological findings could be somehow affected by synaptic effects of these drugs. For example, enhanced NMDAR-mediated signaling due to M1R stimulation by McN-A 343, and thus increased Ca^2+^ entry (Calabresi *et al*., 1998), could interfere with our measurements. However, literature data show that McN-A 343 up to 10 µM (as used here) does not alter SPN membrane properties, which are sensible only to high concentrations (>30 µM) of M1R agonists such as muscarine (Calabresi *et al*., 1998) or carbachol (Hsu *et al*., 1996). On the other hand, it would be interesting to study whether long-term potentiation of corticostriatal synaptic transmission, which depends upon NMDA receptor (Calabresi *et al*., 1992) and M1R activation (Calabresi *et al*., 1999), is altered in *Tshz3*-mutant mice due to their lower cholinergic tone.

In parallel with these experiments, we also assessed whether SPN electrophysiological features in basal conditions (i.e., without drugs) were altered in *Chat-cKO* mice: although average passive (I-V relationship, input resistance and RMP) and active (rheobase and mean firing frequency) membrane properties were unaffected, we observed a drastic change in the distribution of SPN firing frequency values. Namely, instead of a bimodal distribution suggesting two SPN populations with different excitability [D2R-SPNs are more excitable than D1R (Kreitzer & Malenka, 2007; Gertler *et al*., 2008)], *Chat-cKO* mice show a unimodal distribution, suggesting that the two SPN populations are equally or very similarly excitable. Altogether, these electrophysiological findings speak in favor of a lower M1R basal activation by endogenous acetylcholine that differentially alters D1R- and/or D2R-SPN excitability in *Tshz3-*mutant mice. To further this analysis, we used control and *Tshz3-*mutant mice expressing the tdTomato fluorophore in D1R-SPNs (*tdTom-Chat-cKO*), in order to selectively identify and study the two SPN populations. The results obtained with these mice show that the passive membrane properties of D1R- and D2R-SPNs are similar between *tdTom-Chat-cKO* mice and their control, supporting our findings using *Chat-cKO* mice. Moreover, we confirmed literature data showing that, in control mice, D2R-SPNs are more excitable than D1R-SPNs (Kreitzer & Malenka, 2007; Gertler *et al*., 2008), assessed by measuring their rheobase and firing frequency, as well as the D1/D2 ratio of action potential firing frequency per mouse. Interestingly, such D1R- *vs*. D2R-SPN differential excitability is lost in *tdTom-Chat-cKO* mice, where the two SPN populations show similar rheobase and firing frequency, and a D1/D2 ratio close to 1. This could be due to an increased excitability of D1R-SPNs resulting from a reduced basal stimulation of M4Rs by endogenous acetylcholine, without change in D2R-SPN excitability. Indeed, these receptors are mainly expressed by D1R-SPNs and their activation decreases SPN excitability, although such inhibitory effect is not clearly established (Figueroa *et al*., 2002; Hernandez-Flores *et al*., 2015), presumably due to the unavailability of selective M4R agonists. Altogether, our findings in *Tshz3*-mutant mice suggest an unbalanced activity of the direct/indirect pathways, biasing the striatal output in favor of the direct pathway (“go”) rather than the indirect pathway (“no-go”) motor programs, which could contribute to the RBs characterizing this ASD model. A further level of complexity lies on the recent finding that striosomes give rise to both a direct D1 and an indirect D2 pathway targeting substantia nigra pars compacta dopamine neurons that have effects opposite to the canonical matrix-based ones on dopamine release and on movement (Lazaridis *et al*., 2024). This raises the issue of the respective contribution of striosome *vs*. matrix SPNs to the changes reported here, especially since *Chat-cKO* mice exhibit a striosome-biased dysfunction of SCINs (Molitor *et al*., 2026). Moreover, the cholinergic modulation of striatal circuitry is not restricted to M1Rs and M4Rs expressed by SPNs, but also includes the other muscarinic and nicotinic receptors expressed at pre- and postsynaptic level in virtually all striatal neurons types, and on synaptic terminals originating from the cerebral cortex, the thalamus and the substantia nigra pars compacta (Pisani *et al*., 2007; Bonsi *et al*., 2011; Assous, 2021). Therefore, it cannot be excluded that the observed changes could be in part due to other elements of the complex striatal cholinergic system, directly affecting other neurotransmitter systems that modulate SPN excitability. Also, the translational value of our findings obtained using the *Chat-cKO* model is limited, due to the conditional deletion of *Tshz3* in SCINs rather than the global haploinsufficiency found in patients with *TSHZ3* deletion. Nevertheless, these mice still represent a powerful tool to dissect the role of the striatal cholinergic system in RBs associated to ASD.

ASD is a neurodevelopmental disorder for which there are currently no interventions that can improve all symptoms. Based on the data that point to low levels of acetylcholine as potential contributors, treatments based on acetylcholinesterase inhibitors (e.g., donepezil, galantamine and rivastigmine) have been tested in patients (Chez *et al*., 2004; Rossignol & Frye, 2014; Ure *et al*., 2023). However, a recent systematic review found that only 2 out of 680 trials met the criteria required to evaluate the efficacy and adverse effects of these drugs in people with ASD (Ure *et al*., 2023). One of these trials concluded that short-term donepezil treatment may have a limited impact on cognitive functioning in ASD (Handen *et al*., 2011). A more recent review considered 12 studies out of 404 investigating the effect of Alzheimer’s disease medication on neurocognitive outcomes in patients with ASD. Interestingly, 6 of the 12 retained studies reported results following treatment with a cholinesterase inhibitor, 5 of which used donepezil (Diamandis *et al*., 2025). Despite the limited number of studies, the outcome of donepezil treatment for ASD is encouraging, suggesting improvements in language, executive function and complex attention. Overall, these evidences highlight the need for further and conclusive research on the therapeutic effects of this drug on ASD symptoms. Surprisingly, among the studies using mouse models of ASD, only two have examined the impact of donepezil on symptoms related to autism: in BTBR *T^+^ tf/J* mice, donepezil increased social attention, yet failed to improve stereotypical behaviours (Karvat & Kimchi, 2014), while in a valproic acid model it improved social behaviour deficits, RBs and hyperactivity (Kim *et al*., 2014). Here, using a genetic mouse model of ASD (*Chat-cKO*), we demonstrated that acute donepezil treatment significantly reduces MBT score. Interestingly, these findings are consistent with a recent clinical study suggesting that treating autistic patients with donepezil significantly improves stereotypic behaviour (Ghahreman *et al*., 2023). These therapeutic effects of donepezil on RBs clearly support the involvement of reduced cholinergic tone in this ASD-related pathological behavior.

In conclusion, the present study extends our knowledge about the role of the striatal cholinergic system in the pathophysiological mechanisms of RBs associated to ASD. Notably, in a mouse model of ASD characterized by a decreased cholinergic tone in the striatum due to SCIN hypoactivity, we demonstrated that an acute cholinergic treatment by donepezil significantly reduces RBs.

## Data availability statement

Raw data sets will be made available upon reasonable request to the corresponding authors.

## Acknowledgments

We wish to thank: the mouse facility and the “STechBio” service personnel at the CRPN for their helpful technical assistance; Dr. Boris Burle (CRPN) for his kind help with the statistical analysis; Dr. Olivier J. Manzoni and Dr. Pascale Chavis (INMED, INSERM UMR1249, Aix-Marseille Univ, Marseille, France) for kindly providing *Drd1a-tdTomato* mice.

## Author contributions

JG performed the electrophysiological and the behavioral experiments and analyzed the data, under the supervision of PG and AMO. JG and XC generated, maintained and genotyped the transgenic mouse lines. AF performed the RT-qPCR experiments and analyzed the data. LF, XC LK-LG and PG conceived and supervised the project and the experiments. PG wrote the paper with the help and contribution of all the authors. All authors have read and approved the final manuscript.

